# Meta-analysis of genome-wide association studies for height and body mass index in ∼700,000 individuals of European ancestry

**DOI:** 10.1101/274654

**Authors:** Loic Yengo, Julia Sidorenko, Kathryn E. Kemper, Zhili Zheng, Andrew R. Wood, Michael N. Weedon, Timothy M. Frayling, Joel Hirschhorn, Jian Yang, Peter M. Visscher, the GIANT Consortium

## Abstract

Genome-wide association studies (GWAS) stand as powerful experimental designs for identifying DNA variants associated with complex traits and diseases. In the past decade, both the number of such studies and their sample sizes have increased dramatically. Recent GWAS of height and body mass index (BMI) in ∼250,000 European participants have led to the discovery of ∼700 and ∼100 nearly independent SNPs associated with these traits, respectively. Here we combine summary statistics from those two studies with GWAS of height and BMI performed in ∼450,000 UK Biobank participants of European ancestry. Overall, our combined GWAS meta-analysis reaches N∼700,000 individuals and substantially increases the number of GWAS signals associated with these traits. We identified 3,290 and 716 near-independent SNPs associated with height and BMI, respectively (at a revised genome-wide significance threshold of *p*<1 × 10^−8^), including 1,185 height-associated SNPs and 554 BMI-associated SNPs located within loci not previously identified by these two GWAS. The genome-wide significant SNPs explain ∼24.6% of the variance of height and ∼5% of the variance of BMI in an independent sample from the Health and Retirement Study (HRS). Correlations between polygenic scores based upon these SNPs with actual height and BMI in HRS participants were 0.44 and 0.20, respectively. From analyses of integrating GWAS and eQTL data by Summary-data based Mendelian Randomization (SMR), we identified an enrichment of eQTLs amongst lead height and BMI signals, prioritisting 684 and 134 genes, respectively. Our study demonstrates that, as previously predicted, increasing GWAS sample sizes continues to deliver, by discovery of new loci, increasing prediction accuracy and providing additional data to achieve deeper insight into complex trait biology. All summary statistics are made available for follow up studies.

---

Over the past 15 years, genome-wide association studies have been increasingly successful in unveiling many aspects of the genetic architectures of complex traits and diseases^1–6^. GWAS have led to the discovery of tens of thousands of polymorphisms (SNPs in general) associated with inter-individuals differences in quantitative traits or disease susceptibility. They have also been used to generate experimentally testable hypotheses and predict traits and disease risk^7,8^. One of the early challenges faced by GWAS has been to bridge the gap between the amount of trait variance explained by genome-wide significant loci 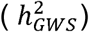 compared to estimates of heritabilities from family-based studies 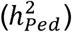. The reasons explaining the gap between 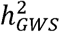 and 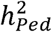, also termed as missing heritability, are now better understood. Contributions from Yang *et al.* (2010; 2015; 2017)^9–11^ and others^12^ have helped clarifying the distinction between what can potentially be explained by all SNPs (a.k.a SNP heritability, 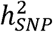) and what remains out of the reach of SNP-array based GWAS, for instance causal variants that are not tagged by genotyped or imputed SNPs. It is worth noting that untagged variants are often rare or even unique to single nuclear families. Therefore, their effects might remain statistically undetectable, despite still contributing to the difference between 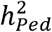 and 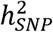. Overall, clarifying the differences between 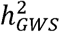, 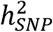 and 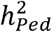 has been a major advance in the field and has helped providing theoretical guarantees that increasing GWAS sample sizes would continue to yield more discoveries, as long as the difference between 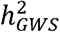 and 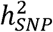 persists.

To date the largest published GWAS of height^5^ and BMI^6^ in ∼250,000 participants on average have uncovered 697 and 97 near-independent SNPs associated with these traits and explaining ∼15% and ∼3% of trait-variance respectively. Compared with 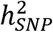 estimates of height and BMI, i.e. ∼50% and ∼30% respectively^9–11^, this indidicates an enormous potential for discoveries expected simply from increasing sample sizes. However, the required sample size to explain all SNP heritability by identified individual genome-wide significant loci is not known because it depends on the joint distribution of allele frequency and effect size at causal variants. Here we perform a meta-analysis of previous GIANT GWAS studies with new GWAS of height and BMI in ∼450,000 participants of the UK Biobank (UKB). In total our sample size reaches ∼700,000 which is unprecedented for GWAS of these traits. The present study is part of a larger effort led by the GIANT consortium, expected in the near future to yield one the largest GWAS ever conducted (*N* between 1.5 and 2 million). We describe below our findings in terms of number of GWAS signals, variance explained, prediction accuracy and also conduct analyses to prioritize genes for follow-up investigations. The summary statistics of these two meta-analyses (height and BMI) are made available (URLs).

## Results

### GWAS of height and BMI identify 3,290 and 716 associated SNPs respectively

We first performed a GWAS of height and BMI in 456,426 UKB participants of European ancestry (Online methods). We tested associations of 16,652,994 genotyped and imputed SNPs (Online methods) with both traits using a linear mixed model to account for relatedness between participants and population stratification. Analyses were performed with BOLT-LMM v2.3^13,14^ using a set of 711,933 HapMap 3 (HM3) SNPs to model the polygenic component to control for relatedness and population stratification (Online methods). After fitting age, sex (inferred from SNP data^15^), 10 genotypic principal components (PCs), recruitment centre, and genotyping batches as fixed effects, phenotypes were residualised (separately for males and females) and inverse-normally transformed before analysis. A parallel analysis of 451,099 individuals and using a slightly different set of parameters for sample selection and adjustment revealed very similar results (Online methods) and so we proceeded with the larger sample of 456,426 UKB participants. We then performed fixed-effect inverse-variance weighted meta-analysis of UKB results with publicly available GWAS summary statistics of height^5^ (GIANT_height_) and BMI^6^ (GIANT_BMI_) using the software METAL^16^. In total, our meta-analysis involves ∼2.4 million HapMap 2 (HM2) SNPs, N = 693,529 participants on average for height and N = 681,275 participants on average for BMI. **Fig. 1** shows Manhattan plots for both meta-analyses. We found in both traits a marked deviation of the distribution of *p*-values from the uniform null distribution (height: λ_GC_ = 3.2; BMI: λ _GC_ = 2.5), suggesting polygenicity and possibly population stratification. The mean of association chi-square statistics is ∼7.2 for height and ∼3.4 for BMI, consistent with a randomly chosen SNP, on average, being associated with height at *p*<0.007 and with BMI at *p*<0.07. We performed LD score regression (LDSC)^17,18^ to quantify the contribution of population stratification to our results. We found that LDSC intercept (I_LDSC_) was inflated for both height (I_LDSC_ = 1.48, s.e. 0.1) and BMI (I_LDSC_ = 1.08, s.e. 0.02), suggesting a significant contribution of population stratification. However, although classically used, this statistic may not accurately reflect the contribution of population stratification as it can rise above 1 with increased sample size and heritability^13^. In contrast, the attenuation ratio statistic R_PS_ = (I_LDSC_ - 1) / (mean of association chi-square statistics-1), which does not have these limitations, was shown to yield a better quantification of population stratification^13^. We found for height and BMI that R_PS_ equals 0.06 (s.e. 0.01) and 0.03 (s.e. 0.01), respectively, which implies that polygenicity is the main driver of the observed inflation of test statistics. We also used the LDSC methodology to estimate the genetic correlation between summary statistics from GIANT_height_ and that from UKB, as well as between summary statistics from GIANT_BMI_ and that from UKB. We found a genetic correlation (*r*_*g*_) of 0.96 (s.e. 0.01) for height and of 0.95 (s.e. 0.01) for BMI, highlighting a strong genetic homogeneity between UKB and previous meta-analyses, and thus confirming the validity of using a fixed-effect meta-analysis. Also, this analysis implied significant overlap of ∼59,000 participants between UKB and GIANT_height_ (bivariate LDSC intercept: 0.17; s.e. 0.05), but not between UKB and GIANT_BMI_ (bivariate LDSC intercept: 0.01; s.e. 0.01). The latter observation is surprising given that the vast majority of cohorts included in GIANT_height_ are also included in GIANT_BMI_. Analogous to the univariate case, we observed in simulations that large sample sizes and heritabilities can inflate the bivariate LDSC intercept even in the absence of sample overlap (**Supplementary Note; Fig S1**). We therefore conclude that sample overlap is likely negligible between UKB and GIANT_height_ as it is between UKB and GIANT_BMI_.

**Fig. 1.**
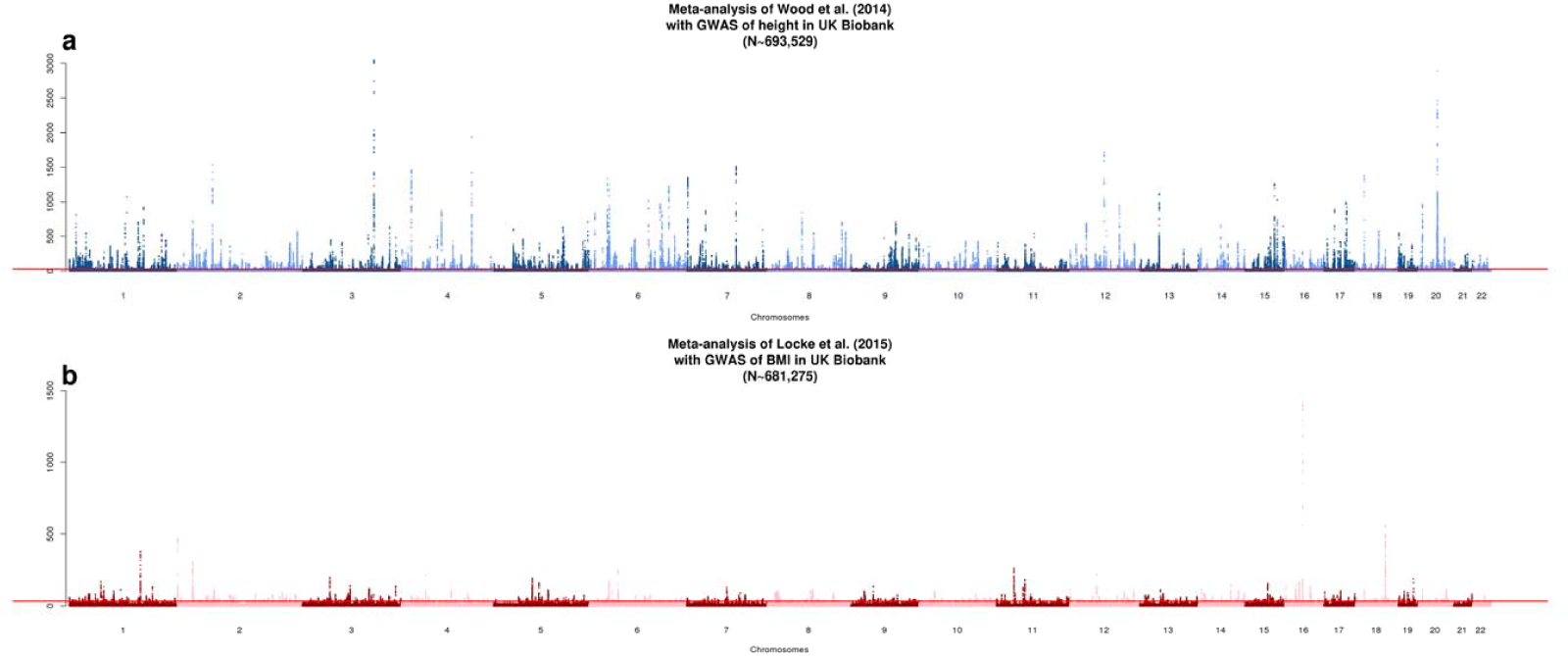
Manhattan plot showing association *χ* ^2^ statistics of association between SNPs and height (panel **a**) or body mass index (BMI, panel **b**).

Using an approximate conditional and joint multiple-SNP (COJO) analysis implemented in GCTA^19^ that takes into account linkage disequilibrium (LD) between SNPs at a given locus, we identified 3,290 and 716 SNPs (COJO *p*<1 × 10^−8^; **Table 1**) associated with height and BMI respectively. This more conservative significance threshold was chosen from the recommendations of a previous study^20^ which showed that type I error was not properly controlled at the classical 5 × 10^−8^ thresold when using SNPs imputed to the Haplotype Reference Consortium or 1,000 genomes imputation reference panels. Compared to GIANT_height_ and GIANT_BMI_, our findings represent a ∼5 and ∼7-fold increase of the number of GWAS signals associated with height and BMI respectively. The 3,290 height-associated SNPs consist of 2,388 primary associations and 902 secondary signals, i.e. genome-wide significant (GWS) in GCTA-COJO analysis only. These 3,290 SNPs clustered in 712 genomic loci (locus is defined as in ref.^5^ as one or multiple jointly associated SNPs located within a 1-Mb window), including 409 loci not previously detected in GIANT_height_. For BMI, the 716 SNPs identified consist of 450 primary associations and 266 secondary signals, clustered in 416 genomic loci including 353 loci not previously detected in GIANT_BMI_. We found that the average number of height and BMI associated SNPs per locus is 4.6 and 1.7 respectively, but also observed a large variability of that number (standard deviation: ∼6 SNPs/locus for height loci and ∼2 SNPs / locus for BMI loci). We found one locus on chromosome 12q23.2 (chr12:102,229,631-103,278,745; genome build hg19) that concentrates up to 19 jointly significant signals for height within ∼1.05 Mb. That locus contains the *IGF1* gene which was previously identified in GIANT_height_. Note however that only 2 independent associations within that locus were reported at that time, indicating that larger GWAS improves the characterisation of the allelic heterogeneity of genomic loci.

**Table 1.** Summary of results from the meta-analysis of GWAS of height and body mass index (BMI) in N∼700,000 individuals of European ancestry and from downstream analyses such as gene-based association tests or Summary-data based Mendelian Randomization (SMR). Prediction accuracy (squared correlation R^2^, between genetic predictors and traits) and variance explained (estimated using GCTA software) is assessed in 8,552 unrelated participants of the Health and Retirement Study (HRS). *New loci refer to loci not identified in Wood *et al*. (2014) or in Locke *et al*. (2015).

| Summary of results | Meta-analysis of height<br>(mean N~693,529) | Meta-analysis of BMI<br>(mean N~681,275) |
| --- | --- | --- |
| Number of genome-wide significant SNPs (GWS; $p < 10^{-8}$) | 3,290 | 716 |
| Number of main / secondary associations | 2,388 / 902 | 450 / 266 |
| Number of loci identified | 712 | 416 |
| Number of new loci* | 409 | 353 |
| Number of genes identified through SMR analysis | 610 | 110 |
| Number of methylation sites identified through SMR analysis | 775 | 176 |
| Prediction accuracy ($R^2$) in HRS from GWS SNPs | 19.7% | 4.1% |
| Prediction accuracy ($R^2$) in HRS from SNPs at $p < 0.001$ | 24.4% | 8.6% |
| Variance explained in HRS from GWS SNPs | 24.6% | 5.0% |
| Variance explained in HRS from SNPs at $p < 0.001$ | 34.7% | 10.2% |

We assessed the replicability of these associations by estimating the regression slope of SNP effect size estimated in an independent sample onto the SNP effect sizes (corrected for winnner’s curse effects^21,22^) from our meta-analyses, using 8,552 unrelated individuals from the Health and Retirement Study (HRS). A similar approach to quantify replicability was been applied in Turley et al. (2017)^23^. We found significant regression slopes for height (0.90; s.e. 0.02) and BMI (0.91; s.e. 0.04), both close to one and therefore suggesting a high level of replicability of our findings (**Fig. 2**).

**Fig. 2.**
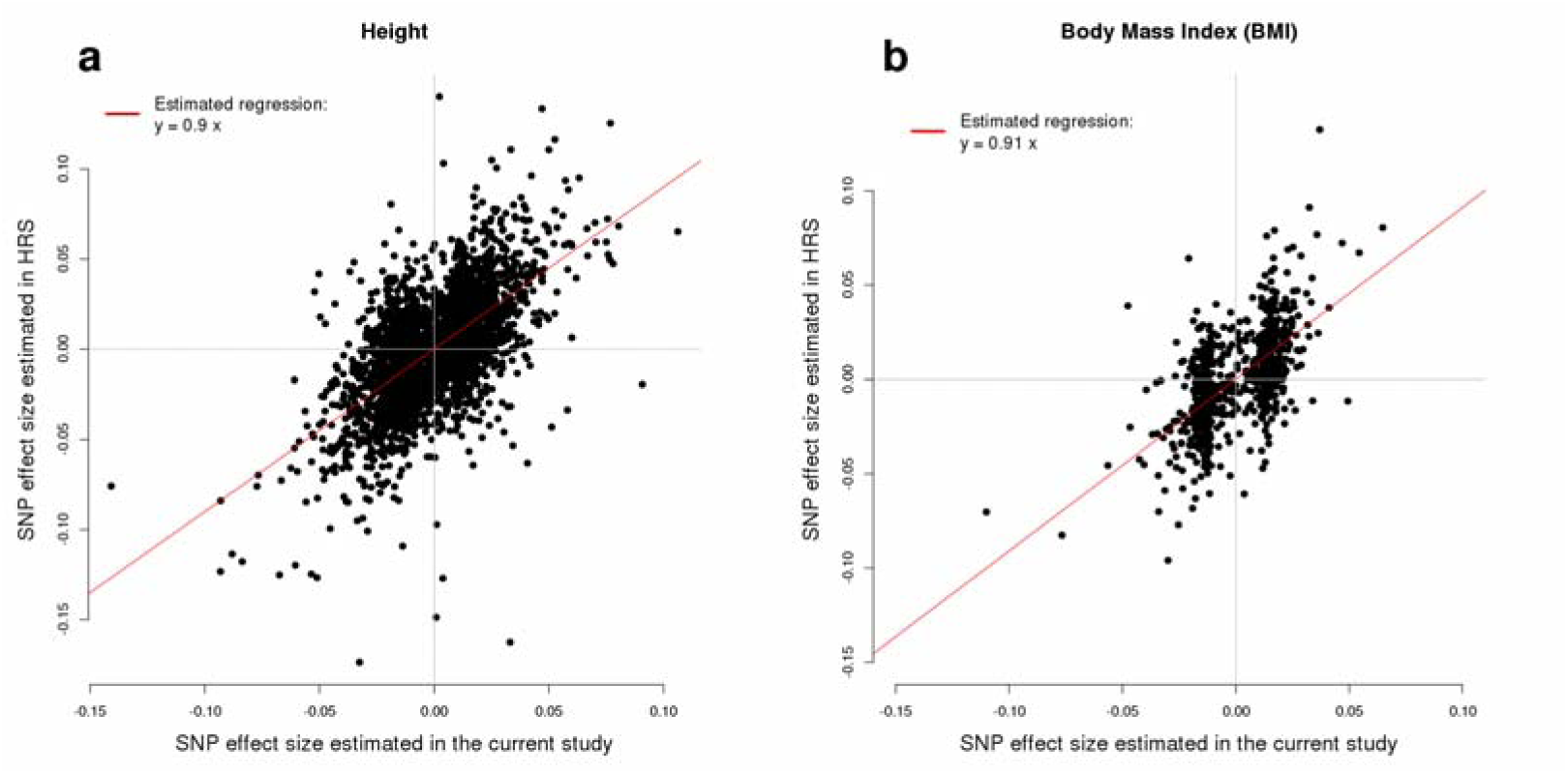
Regression of SNPs effect estimated from the meta-analysis of GWAS of height in UKB and GWAS of height from Wood et al. (2014) (panel **a**); and GWAS of body mass index (BMI) in UKB and GWAS of BMI from Locke et al. (2015) (panel **b**) onto SNP effects on height and BMI estimated in HRS.

### Predictive power of polygenic scores

We estimated in HRS participants using the GCTA-GREML approach^9,10,19^ that GWS SNPs explain 24.6% (s.e. 1.3%) and 5.0% (s.e. 0.7%) of the variance of height and BMI, respectively, adjusting for 20 PCs for both traits. This represents a ∼1.5 and ∼2.6-fold improvement in comparison with previous meta-analyses (**Fig. 3a-b**). For each HRS participant, we also calculated genetic predictors of height and BMI from GWS SNPs as the sum of trait increasing alleles at these loci, weighted by their estimated effect sizes. We found the squared correlation between predicted height and actual height to be ∼19.7% and between predicted BMI and actual BMI to be ∼4.1% (**Fig. 3c-d**). We performed additional prediction analyses using SNPs with significance *p*-values larger than 10^−8^. We performed GCTA-COJO analyses for height and BMI and analysed SNPs with significance *p*-value below 10^−3^, 10^−4^, 10^−5^, 10^−6^, 10^−7^ and 10^−8^. We therefore calculated 6 genetic predictors for each trait and quantified in HRS participants the fraction of trait variance explained by SNPs contributing to these predictors as well as their predictive capacity (**Fig. 3**). As reported in Wood *et al*. (2014), we found that including SNPs beyond GWS increases prediction accuracy and variance explained (**Fig. 3**) in both traits. For height, the variance explained increased from ∼24.6% using 3,290 GWS SNPs to ∼34.7% (s.e. 1.9%) using ∼15,000 SNP with *p*<0.001. The prediction *R*^2^ also increased from ∼19.7% to ∼24.4%. For BMI, the variance explained using ∼9000 SNPs selected in the COJO analysis at *p*<0.001 is ∼10.3% (s.e. 1.4%) and the corresponding prediction *R*^2^ is ∼8.6%, which is twice the prediction accuracy obtained using GWS loci only.

**Fig. 3.**
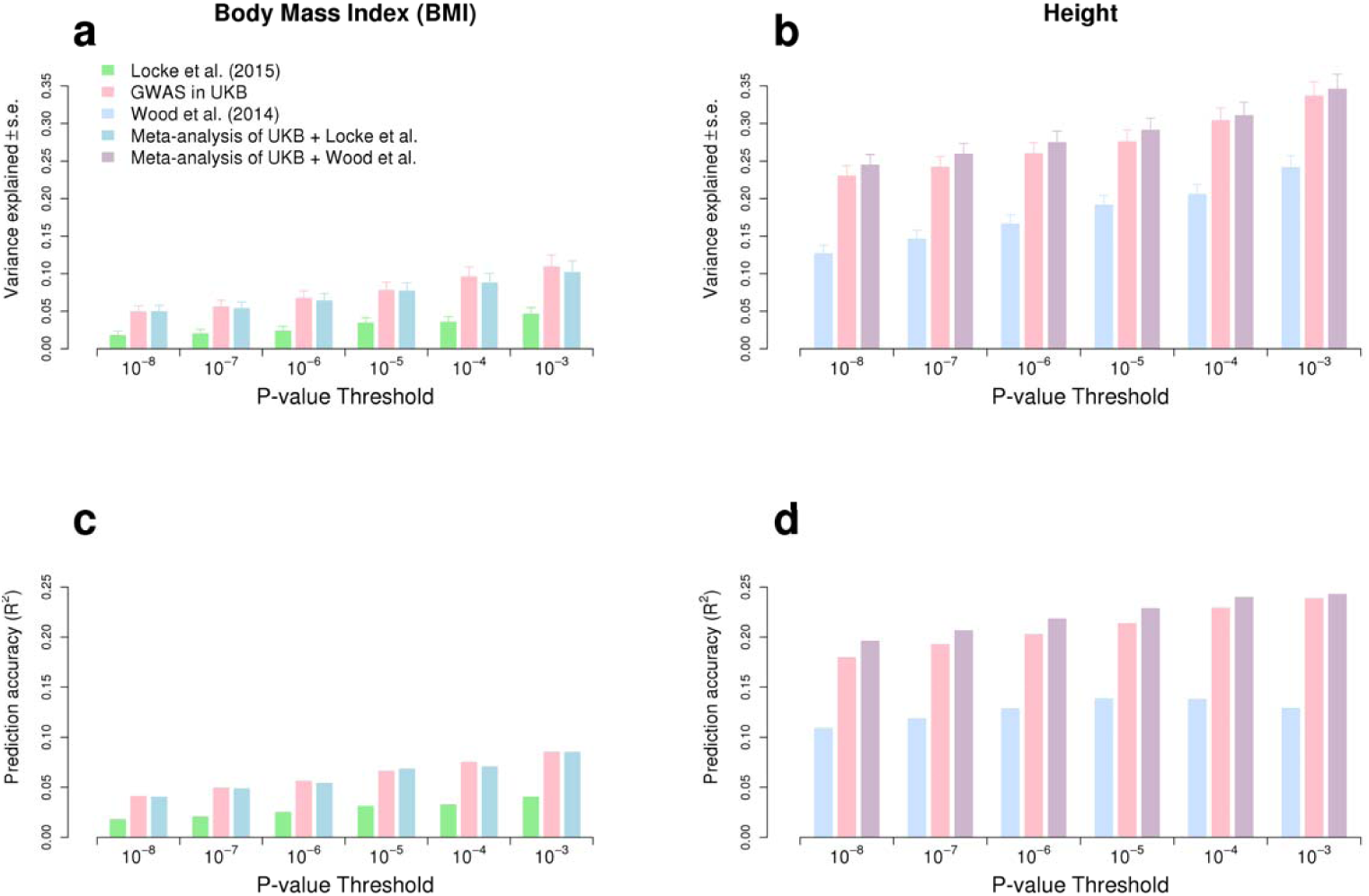
Variance explained and prediction accuracy (squared correlation between trait value and its predictor from SNPs) calculated from 6 nested sets of SNPs selected at different significance threshold. Variance explained and prediction accuracy is calculated among 8,552 unrelated participants of the HRS cohort.

### Gene prioritization

We next attempted to identify genes whose expression levels could potentially mediate the association between SNPs and height or between SNPs and BMI. For this purpose, we performed a summary-data based Mendelian randomization (SMR) analysis^24^. This method aims at testing the association between gene expression (in a particular tissue) and a trait using the top associated expression quantitative trait loci (eQTL) as a genetic instrument. For this analysis, which only requires GWAS summary statistics, we used the publicly available GTEx-v7 database containing eQTLs for multiple genes in multiple tissues^25^. We identified 610 (**Table 1**; including 444 identified by gene-based test) and 110 (**Table 1**; including 74 identified by gene-based test) unique genes which genetic control suggestively overlaps (*p*_SMR_<5 × 10^−8^) that of height or BMI. Significant SMR test indicates evidence of causality or pleiotropy but also the possibility that SNPs controlling gene expression are in linkage disequilibrium with those associated with the traits. These two situations can be disentangled using the HEIDI (HEterogeneity In Dependent Instrument) test implemented in the SMR software. The number of genes reported above corresponds to genes already filtered on statistical evidence supporting pleiotropy or causality rather than linkage between variants controlling gene expression and variants controlling height or BMI (*P*_HEIDI_ > 0.05). We found that >95% (597/610 = ∼98% for height and 105/110 = ∼95% for BMI) of height-and BMI-associated genes identified via the SMR analysis show consistent direction of effects across multiple tissues. As an example, we found that higher expression of *PIGP* across 23 tissues is associated with increased height; and that higher expression of *HSD17B12* across 41 tissues is associated with decreased BMI. We then quantified the enrichment of the genes identified via SMR and HEIDI tests, into biological pathways. Altogether, we found that height-associated genes are significantly enriched among genes contributing to skeletal growth, cartilage and connective tissue development; while BMI-associated genes are mostly enriched among genes involved in neurogenesis and more generally involved in the development of the central nervous system. These last results therefore confirm findings from Wood *et al*. (2014) and Locke *et al*. (2015) which previously implicated the same pathways and highlighted their connections with height and BMI.

### Mediation through epigenetic mechanisms

We performed another SMR and HEIDI analysis to now prioritise CpG dinucleotides which methylation levels mediate the association between SNPs and height or BMI. For this analysis, we used publicly available methylation QTL (*m*QTL) from the McRae *et al*. (2017) study^26^ in peripheral blood. We identified 775 and 176 (**Table 1**) DNA methylation sites showing pleiotropic associations with height and BMI respectively. Among all CpG sites identified, we found that increased DNA methylation at *cg19825988* (within the *ZBTB38* gene) has the largest positive mediation effect on height (∼0.4 SD for 100% methylation; *p*_SMR_ = 3.5 × 10^−9^). The *ZBTB38* gene, located within a previously identified GWAS locus (GIANT_height_), encodes a zinc finger transcriptional activator that binds methylated DNA. This gene was also detected in our first SMR analysis (using gene expression) described above. For BMI, the largest effect of DNA methylation was observed at *cg03755535* (within the first exon of the *CAMKV* gene); where decreased DNA methylation correlates positively with increased BMI (- 0.14 SD for 100% methylation; *p*_SMR_ = 4.9 × 10^−8^). This gene was not detected in our first SMR analysis but is located within a previously identified GWAS locus.

## Discussion

We have presented here the results of the meta-analysis of a single large study, the UK Biobank, with previously published GWAS of height and BMI. We found that the number of genomic loci associated with height and BMI is disproportionately increased compared to previously published GWAS, and that this increase correlates with increased trait variance explained and improved accuracy of genetic predictors from SNPs at these loci. In addition, we have shown that large GWAS enhance the power of integrative analyses such as pathway enrichment and summary-data based Mendelian randomization to unveil relevant genes to be prioritized for further functional studies.

Our analyses revealed a number challenges to address when dealing with very large GWAS. One of these challenges relates to conclusions from LDSC, a method now routinely used for quality control (detection of confounding effects) and inference of genetic parameters like heritability and genetic correlation. Following the recent study by Loh *et al*. (2018), which pointed out a number of caveats relative to the interpretation of the univariate LDSC intercept as an indicator of confounding due to population stratification or other artefacts, we have shown here that caution must also be applied when interpreting the intercept of the bivariate LDSC. These two problems, which are directly related to each other, both illustrate how the effect of very subtle population stratification can be dramatically magnified when sample sizes are large. We recall here the suprising observation that, despite considering the same sets of cohorts, the conclusions about sample overlap from bivariate LDSC intercept were radically different between GWAS of height and GWAS of BMI. Similar to the univariate case, we recommend the use of an attenuation ratio statistic to measure how much of the inflation in the bivariate LDSC intercept is explained by correlated population stratification or sample overlap.

Another challenge faced in this study relates to the over-correction of population stratification. In general, setting up expectations with respect to how many GWAS signals can be reasonably detected or how much variance can be explained from SNPs identified in a GWAS of a given sample has always been a difficult question. In particular, the detection of “too many” GWAS signals has often been a concern in the GWAS literature and seen as an indication of potentially uncorrected population stratification. With very large datasets like UKB, some of these questions can be now addressed. We observed that the number of variants and fraction of variance explained by GWAS hits identified from random subsets of UKB of the same size as GIANT_height_ or GIANT_BMI_ was larger than that discovered in those studies (**Table 2**). Multiple reasons could explain these differences, as for example genetic and phenotypic heterogeneity between cohorts included in these two meta-analyses^27^. Nonetheless, we argue that methods classically used to correct for the effects of population stratification may have removed a substantial amount of the signal to be detected. To further illustrate this point, we re-analysed the data from Locke *et al*. (2015). Our new analysis consisted of deflating the genomic control (GC) corrected standard errors of estimated SNP effects with a factor equal to the square-root of the LDSC intercept. This transformation constrains the LDSC intercept to be 1 but is less conservative than the double GC correction (i.e. GC correction in each cohort included in the meta-analysis, and GC correction applied to the outcome of the meta-analysis) used in Locke *et al*. (2015). We found in this secondary analysis that the number of GWAS signals (at *p*<10^−8^) increased from 77 to 210 (∼3-fold increase), the variance explained increased from ∼1.8% to ∼2.8% and the prediction accuracy of genetic predictors using those SNPs from 1.8% to 2.4% (**Fig. S2**). This observation demonstrates that a correction based on LDSC intercept performs better than GC correction but still remains imperfect, since we know that LDSC intercept also increases with sample size. New methods must therefore be developed in order to maximize the potential of discovery of forthcoming GWAS of ever-larger sample sizes.

**Table 2.** Number, percentage of variance explained and accuracy of genetic predictors from SNPs found associated (*p*<10^−8^) with height or body mass index (BMI) in a random sample of 250,000 unrelated participants of the UKB. For comparison, similar statistics are reported from GWAS hits identified in Wood *et al*. (2014) and Locke *et al*. (2015). Prediction accuracy (squared correlation R^2^, between genetic predictors and traits) and variance explained (estimated using GCTA software) is assessed in 8,552 unrelated participants of the Health and Retirement Study (HRS).

|  | Height |  | BMI |  |
| --- | --- | --- | --- | --- |
|  | Random sample<br>from UKB | Wood et al. (2014) | Random sample<br>from UKB | Locke et al. (2015) |
| Number of genome-wide<br>significant SNPs (GWS; $p < 10^{-8}$) | 850 | 594 | 160 | 82 |
| Variance explained by GWS | 14.0% | 12.8% | 2.3% | 1.8% |
| Prediction accuracy ($R^2$) | 14.0% | 10.9% | 2.5% | 1.8% |

In summary, our study confirms the potential for new discoveries of large genome-wide association studies and announces a gargantuan number of new discoveries for the next iteration of meta-analyses of the GIANT consortium based on sample sizes in the order of 1 million and more.

## Online methods

### UK Biobank analyses

#### Sample selection

We analysed data from 488,377 genotyped participants of the UK Biobank (UKB). We restricted the analysis to 456,426 participants of European ancestry. Ancestry was inferred using a two-stage approach. The first step consisted of projecting each study participant onto the first two genotypic principal components (PC) calculated from HapMap 3 SNPs genotyped in 2,504 participants of 1,000 genomes project^28^. We then used five super-populations (European, African, East-Asian, South-Asian and Admixed) as reference and assigned each participant to the closest population. Distance was defined as the posterior probability under a bivariate Gaussian distribution of each participant to belong to one of the five super-populations. This method generalizes the *k*-means method and takes into account the orientation of the reference cluster to improve the clustering. Vectors of means and 2 × 2 variance-covariance matrices were calculated for each super-population, using a uniform prior.

#### SNP selection

We analysed SNPs imputed to the Haplotype Reference Consortium (HRC) imputation reference panel^29^ with an imputation quality score above 0.3. For each UKB participant, we hard-called genotypes with posterior probability larger than 0.9 and kept SNPs with call rate >0.95, minor allele frequency >0.0001, and *p*-value for Hardy-Weinberg test larger than 10^−6^. In total we analysed 16,653,239 SNPs. For the meta-analysis, we considered a subset of ∼2.3 millions (out of 16,653,239) SNPs showing consistent alleles with UKB and HRS (used as LD reference) as well as consistent allele frequency (maximum difference < 0.15 for minor and major allele).

#### Association testing

We ran a genome-wide association study of height and body mass index (BMI) in 456,426 UKB participants using linear mixed model association testing implemented in BOLT-LMM v2.3^13^ software assuming an infinitesimal model. We used 657,524 HapMap 3 SNPs (LD pruned for SNPs with r^2^ > 0.9) as model SNPs in our analysis. Height and BMI were adjusted for age, sex, recruitment centre, genotyping batches and 10 PCs calculated from 132,102 out the 147,604 genotyped SNPs pre-selected by the UK Biobank quality control team^15^ for principal component analysis. The difference is explained by the quality control of SNPs (minor allele frequency >0.01, genotype call rate > 95% and Hardy-Weinberg test *p*-value > 10^−6^) applied to a different set of samples as compared to Ref^15^. PCs were calculated using the flashPCA software^30^. In a parallel effort, to provide a sensitivity analysis, using slightly different parameters for sample selection and adjustment, we selected 451,099 indivdiuals of European genetic ancestry as described in Ref.^31^. Analyses performed on this second set of UKB participants revealed highly concordant findings compared with the set of 456,426 participants. We therefore reported here results from the larger set of individuals.

### Replication

We used genotypes imputed to the 1,000 genomes reference panel and phenotypes (height and body mass index) from 8,552 unrelated (GRM < 0.05) participants of the Health and Retirement Study (HRS) to assess the replicability of SNPs found to be associatd with height and BMI. We also used these data to assess the variance explained by different sets of SNPs as well as the out-of-sample prediction accuracy of genetic predictors using these sets of SNPs. Analyses were restriced to 2,484,330 HapMap 2 SNPs with an imputation quality score >0.3, a minor allele frequency >0.01 and a *p*-value from Hardy-Weinberg equilibrium test >10^−6^.

Given that replication of individual SNP is not feasible because of the limited sample size of our replication cohort, we assessed the overall replicability of SNP-traits associations using the regression slope of estimated SNP effects from the replication study onto estimated SNP effect sizes from the discovery study. Values of this slope of ∼1 indicate good replicability of GWAS findings. SNPs brought forward for replication are subjected to the winner’s curse effect and their effect sizes are biased^21,22^. We therefore used the correction proposed by Zhong & Prentice (2018) (ref.^20^) before estimating the replication slope.

### Summary statistics QC and meta-analyses

Summary statistics of GWAS of height and BMI from the Wood *et al*. (2014) and Locke *et al*. (2015) studies were downloaded from the following website: https://portals.broadinstitute.org/collaboration/giant/index.php/GIANT_consortium_data_files. Before meta-analysis with UKB, we filtered out SNPs which reported pairs of alleles did not match the pairs of alleles in the HRS and UKB and also which had reported allele frequency too different (absolute difference > 0.15) from that calculated using unrelated participants of HRS. Fixed-effect inverse variance weighted meta-analysis was performed using the software METAL^16^.

### Linkage disequilibrium score regression

We performed linkage disequilibrium (LD) score regression to quantify the level of confounding in GWAS due to population stratification as well as quantifying the sample overlap between cohorts involved in previous meta-analyses and the UK Biobank. Analyses were performed using the LDSC software v1.0.0 (https://github.com/bulik/ldsc). We used default parameters but did not apply any threshold on the maximum association chi-square statistics of SNPs included in the analyses. We used LD scores from Europeans participants of the 1,000 genomes project that can be downloaded from the LDSC website.

### SMR and HEIDI analyses

Summary-data based Mendelian Randomization (SMR) and HEterogeneity In Dependent Instrument (HEIDI) tests were implemented in the SMR software (http://cnsgenomics.com/software/smr/). SMR analyses were performed using default parameters but specifying a window of 2 Mb up- and downstream genes (expression probes) to include relevant cis-eQTL (instrument) for those genes. The same approach was applied for detecting CpG methylation sites associated with height or BMI. SMR analyses were based on eQTLs from publicly available databases from GTEx-v7^25^ and McRae *et al*. (2017) ^26^. Both sets of eQTL in SMR format can be downloaded from the SMR website: http://cnsgenomics.com/software/smr/.

### Data download

GWAS summary statistics can be downloaded from the GIANT consortium website: https://portals.broadinstitute.org/collaboration/giant/index.php/GIANT_consortium_data_files.

## Acknlowledgements

This research was supported by the Australian Research Council (DP130102666, DP160103860, DP160102400), the Australian National Health and Medical Research Council (1078037, 1078901, 1103418, 1107258, 1127440 and 1113400), the National Institute of Health (NIH grants R01AG042568, P01GM099568 and R01MH100141), the Sylvia & Charles Viertel Charitable Foundation. We used genotypic genotypic and phenotypic data from the Health and Retirement Study (HRS: dbGaP phs000428.v1.p1), as well as genotypic and phenotypic data from the UK Biobank under projects 12505 and 12514. We also used eQTL data from The Genotype-Tissue Expression (GTEx) Project, which was supported by the Common Fund of the Office of the Director of the National Institutes of Health, and by NCI, NHGRI, NHLBI, NIDA, NIMH, and NINDS. The GTEX data used for the analyses described in this manuscript were obtained on dbGaP accession number phs000424.v6.p1.

## Supplementary Note

We performed a simulation to quantify the inflation of the bivariate LD score regression (LDSC) intercept created when the sample size of each GWAS and the heritability are large. We used for our simulations genotypes at 1,123,348 HapMap 3 SNPs (Online methods) from 348,502 unrelated (genetic relationship < 0.05) participants of the UK Biobank (UKB) with European ancestry (Online methods). To mimic independent GWAS, we randomly split our dataset in two sub-samples of equal size (N_1_ = N_2_ = 174,251), and simulated 9 traits with the same 10,000 causal variants (randomly sampled among HapMap 3 SNPs) and with an heritability varying from 0.1, 0.2,…, up to 0.9. Each trait was simulated with same SNPs effect sizes in each sub-sample so that the genetic correlation is expected to be 1. We then performed a GWAS of these nine simulated traits in each sub-sample separately, then used GWAS summary statistics to perform a bivariate LD score regression. LD score regression was performed using the LDSC softare v1.0.0 and using LD scores from European samples of the 1,000 genomes reference panel. We present the results our this simulation in **Fig. S1**. Overall, we found when the heritability is larger than 0.5, the bivariate LDSC intercept can be as large as ∼0.1 (s.e. 0.02), which would falsely indicate a potential overlap of 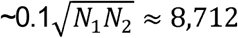 samples between the two sub-sets. We also observed an inflation of the univariate LDSC intecept (Fig. S1, panel **b**), but as mentioned previously, this observation is expected under the theory derived in Bulik-Sullivan *et al*. (2014).

**Fig. S1.**
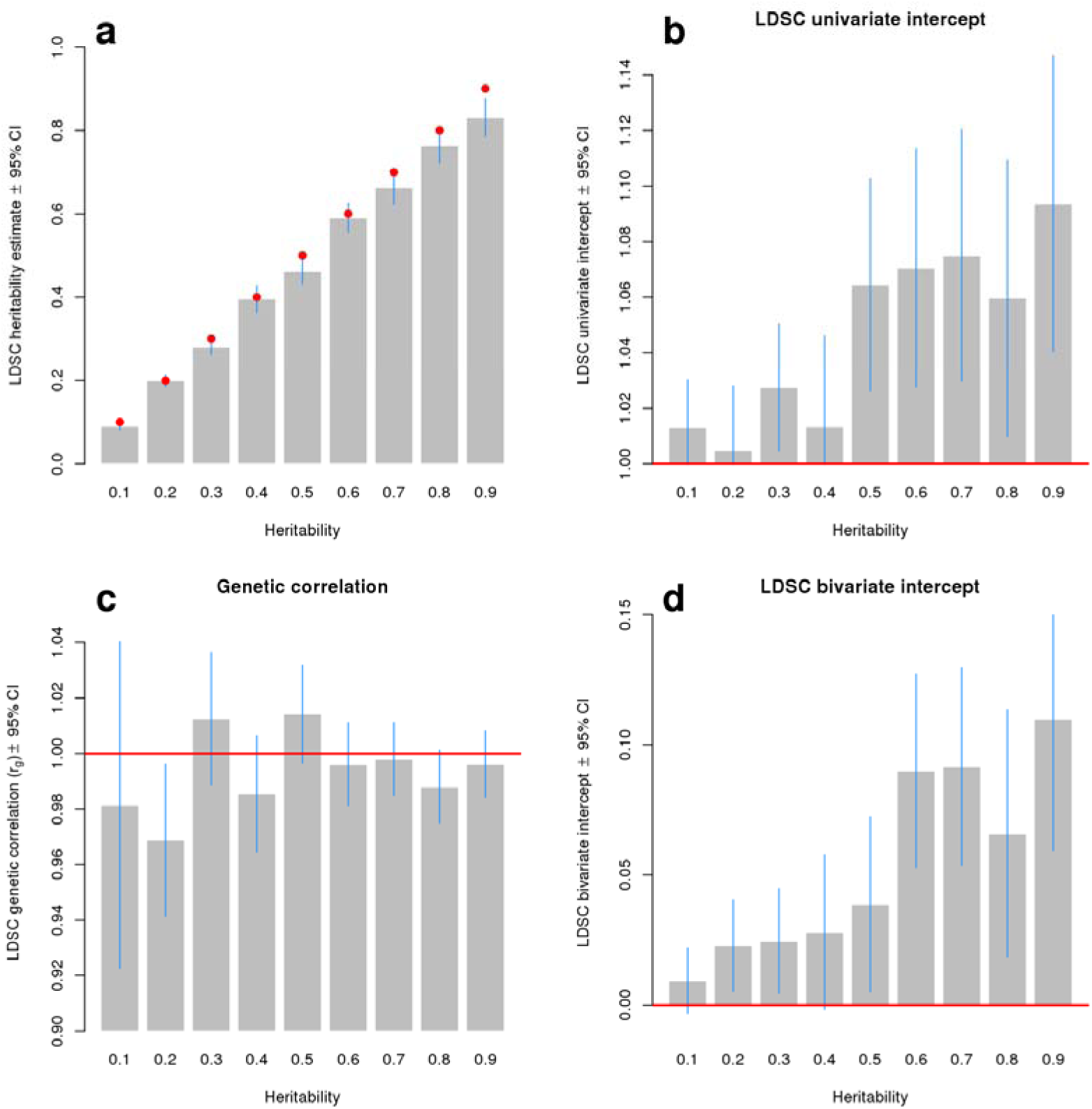
Statistics from the LD score regression applied to 9 simulated GWAS in two independent (non-overlaping) sub-samples of unrelated participants of the UK Biobank (**Supplementary Note**). Panels **a** and **b** show univariate LD score regression intercepts and estimates of heritability respectively obtained from analyzing summary statistics from each sub-sample separately, then averaged between the two independent sub-samples. Panels **c** and **d** show estimates of genetic correlations (expected to be equal to 1, **Supplementay Note**) between the two sub-samples and bivariate LD score regression intercepts respectively, indicating sample overlap between the two sub-samples of UKB, in particular when the underlying heritability is > 0.5. The latter observation illustrates that biviate LD score regression intercept can be inflated even in the absence of sample overlap.

**Fig. S2.**
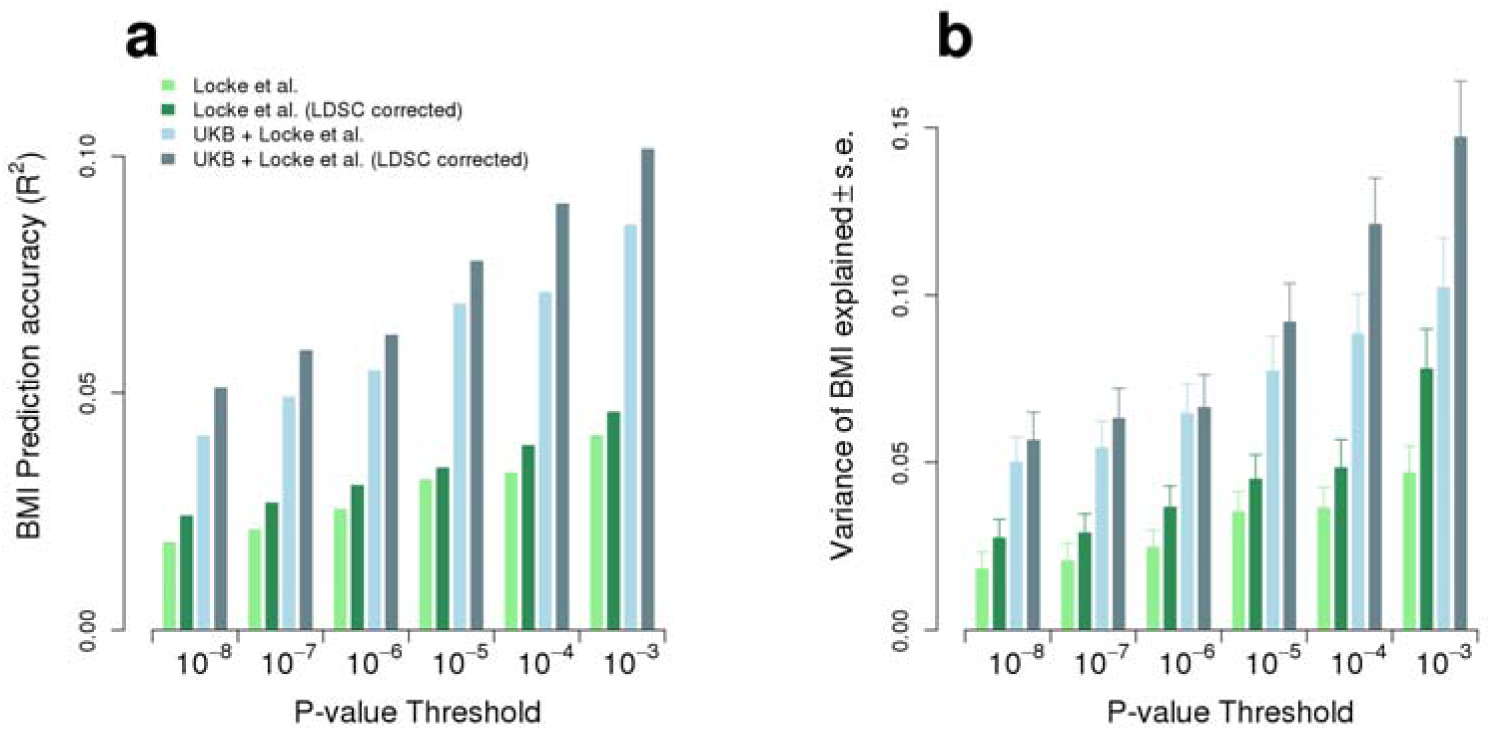
Variance explained and prediction accuracy (squared correlation between the trait value and SNP preditors of that trait) of genetic predictors calculated from 6 nested sets of SNPs selected at different significance threshold. Variance explained and prediction accuracy is calculated among 8,552 unrelated participants of the HRS cohort. Two versions of the GWAS summary statistics from the Locke *et al*. (2015) study were used: Locke et al., as released from the initial publication and Locke et al. (LDSC corrected) in which test statistics were inflated with the inverse of the LD score intercept (here I_LDSC_ = 0.68). The latter method increases power while constraining the LD score intercept to be exactly 1. These two sets of summary statistics were meta-analysed with GWAS summary statistics from UK Biobank (UKB) and resulting asscocation statistics are labeled UKB + Locke et al. and Locke et al. (LDSC corrected) respectively.

